# A Scalable, Easy-to-Deploy, Protocol for Cas13-Based Detection of SARS-CoV-2 Genetic Material

**DOI:** 10.1101/2020.04.20.052159

**Authors:** Jennifer N. Rauch, Eric Valois, Sabrina C. Solley, Friederike Braig, Ryan S. Lach, Morgane Audouard, Jose Carlos Ponce-Rojas, Michael S. Costello, Naomi J. Baxter, Kenneth S. Kosik, Carolina Arias, Diego Acosta-Alvear, Maxwell Z. Wilson

**Author notes:** These authors contributed equally to this work (Jennifer N. Rauch, Eric Valois). These authors are co-senior authors (Diego Acosta-Alvear, Maxwell Z. Wilson).

## Abstract

The COVID-19 pandemic has created massive demand for widespread, distributed tools for detecting SARS-CoV-2 genetic material. The hurdles to scalable testing include reagent and instrument accessibility, availability of highly-trained personnel, and large upfront investment. Here we showcase an orthogonal pipeline we call CREST (Cas13-based, Rugged, Equitable, Scalable Testing) that addresses some of these hurdles. Specifically, CREST pairs commonplace and reliable biochemical methods (PCR) with low-cost instrumentation, without sacrificing detection sensitivity. By taking advantage of simple fluorescence visualizers, CREST allows for a binary interpretation of results. CREST may provide a point-of-care solution to increase the distribution of COVID-19 surveillance.

## Introduction

The COVID-19 pandemic presents the world with an unprecedented public health challenge. The lack of COVID-19 symptoms in a significant proportion (estimates range from 18 to 29%) of SARS-CoV-2 -infected individuals fuels covert transmission of the virus^1,2^. Even in cases in which symptoms do present, the virus can be transmitted before symptom onset^3,4^. The high proportion of asymptomatic, mildly symptomatic, and highly infectious individuals, combined with the risk of severe disease outcomes, and mortality rates ranging from 1.38% to as high as 20% in people 80 years and older, places an unprecedented burden on our healthcare system^5,6^.

A range of mitigation efforts have been implemented across the globe to slow the transmission and “flatten the curve,” from sheltering-in-place to the requirement for face-covering in public. However, without widely-available and accurate testing, it remains unclear exactly how effective these methods are at curbing the spread of the virus. In addition, these risk mitigation efforts have a catastrophic economic and social impact, which makes long-term compliance challenging until a vaccine or treatment is widely available^7^. To reduce the socioeconomic burden, we need to implement effective containment measures, such as contact tracing, quarantine of confirmed cases, and enhanced surveillance, all of which depend on data obtained from widespread testing^8^. Proactive, prevalent, and regular testing have been effective measures for the control of COVID-19 in countries such as South Korea and Taiwan, and will be required for the establishment of effective strategies for the relaxation of social-distancing measures worldwide^9,10^.

A critical hurdle to deploying massively widespread, recurrent testing is the availability of reagents and specialized equipment. Suggested solutions to make sample collection scalable include laboratory-made Viral Transport Media and 3D-printed swabs and bypassing biochemical RNA extraction through heat and chemical extraction techniques^11-14^. Yet, there have been few end-to-end solutions to SARS-CoV-2 testing that fulfill the requirements of being immediately scalable and low-cost, without sacrificing sensitivity. Here, we focused on developing a CRISPR-based SARS-CoV-2 RNA detection method that is low cost, highly sensitive, and easy to deploy at sites with minimal infrastructure.

Cas12 and Cas13 (CRISPR-based) methods have been transformative with regards to pathogen detection. They are sensitive and, when coupled to isothermal amplification methods and lateral flow immunochromatography detection, have been made field deployable. These methods are promising tools for the detection of SARS-CoV-2^15–18^. Yet, because of the global demand for testing, key reagents in these protocols are difficult to obtain. To lower the barrier to COVID-19 diagnostics, we devised a method we call CREST (Cas13-based, Rugged, Equitable, Scalable Testing). CREST addresses three of the main hurdles—reagent accessibility, equipment availability, and cost—that limit the scalability of Cas13-based testing, by taking advantage of widely available enzymes, low-cost thermocyclers, and easy-to-use fluorescent visualizers. Moreover, CREST is equivalent in sensitivity to the gold standard reverse transcription quantitative polymerase chain reaction (RT-qPCR) method most often deployed for COVID-19 testing. With these advantages, CREST has the potential to facilitate early detection of positive cases, regular monitoring of individuals at high risk, and implementation of informed containment measures for infected individuals.

## Results

To design a sensitive, low-cost, and easy-to-use SARS-CoV-2 detection method, we first identified critical steps that require limiting reagents, specific equipment, and highly-trained individuals to perform them, and thus present a barrier to testing at sites with limited infrastructure and resources. For CRISPR-Cas13-based methods, these steps include (i) the amplification of the target material prior to detection, and (ii) the visualization of Cas13 activity either by colorimetric (immunochromatography) or fluorescent methods.

To lower the first barrier, we analyzed options for detection of specific SARS-CoV-2 genomic sequences upon enzymatic amplification (Fig. 1). The gold standard method relies on RT-qPCR^19^. Quantitative detection is accomplished using specialized instruments that detect fluorescent probes which report the extent of amplification of the target sequence in real time. While sensitivity is high (on the order of tens of target molecules per microliter), the main limitation is the requirement of real-time thermocyclers, analysis software, and trained personnel for data interpretation. Limitations aside, the core of the technology—amplification of a target nucleic acid sequence by PCR—is robust, sensitive, and makes use of a widely-available enzyme, Taq polymerase. These advantages motivated us to pair PCR with CRISPR-based detection of viral sequences, an approach that has been previously successful for the detection of DNA sequences using Cas9^20^. The thermocyclers required for PCR are expensive, specialized instruments that are generally limited to professional laboratories. However, the recent DIY-Bio movement has made PCR accessible through the creation of affordable, Bluetooth-enabled, field-ready thermocyclers, which can even be battery operated. These versatile thermocyclers can be used in unconventional environments and perform as well as traditional thermocyclers in moderate temperatures^21–23^. We reasoned that these devices, such as the miniPCR mini16, offer a low-cost solution for the amplification of the viral target material and can make COVID-19 testing widely available (Fig. 2).

**Figure 1.**
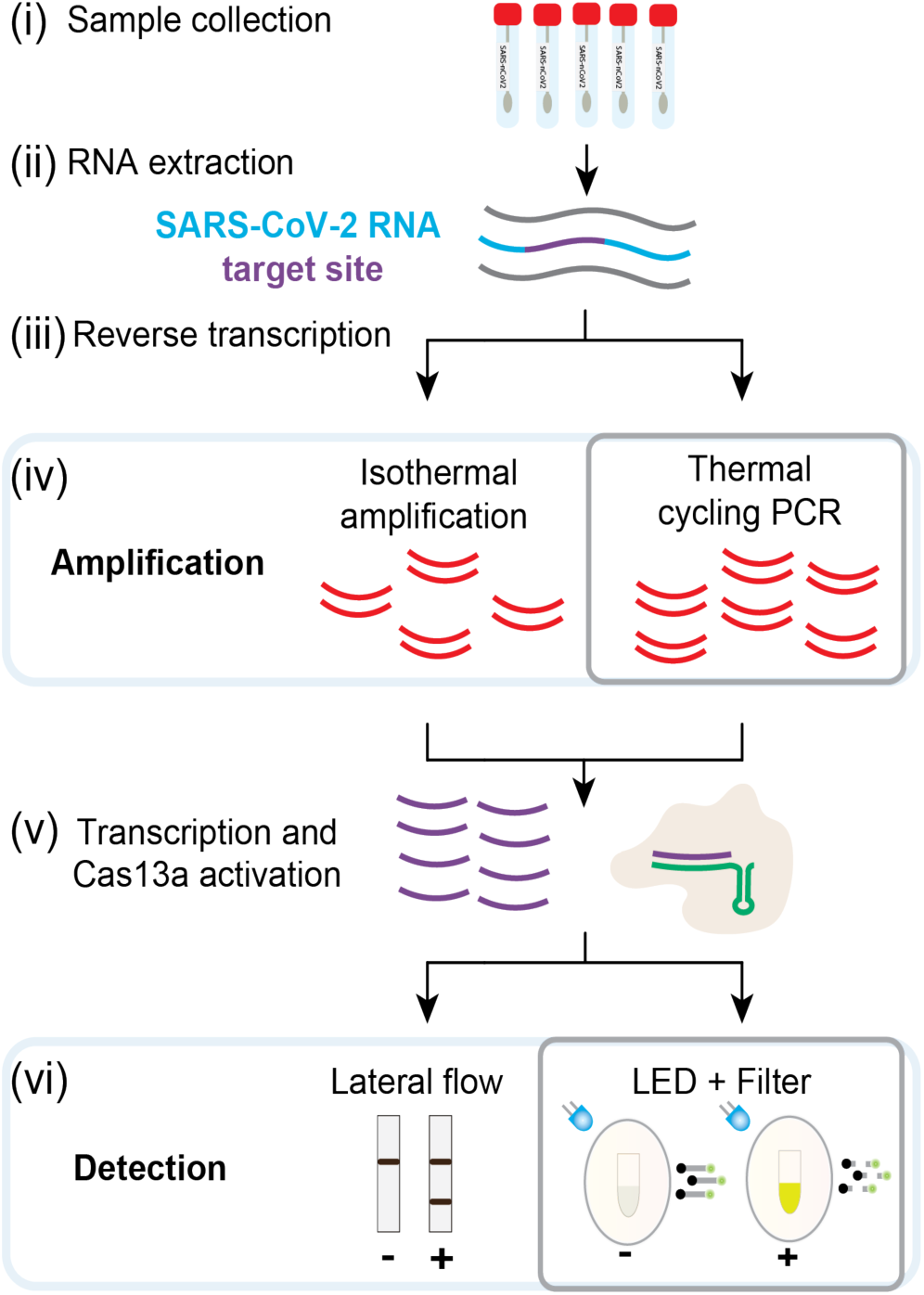
Overview of Cas13-based detection methods and CREST modifications. (i-iii) Standard sample collection, RNA extraction, and reverse transcription. (iv) Amplification using cost-effective Taq polymerase and portable thermocyclers instead of isothermal reactions. (v) Transcription and Cas13 activation are followed by fluorescence detection of de-quenched poly-U cleavage reporter visualized with a blue LED (∼495 nm) and orange filter or other fluorescence detection system.

**Figure 2.**
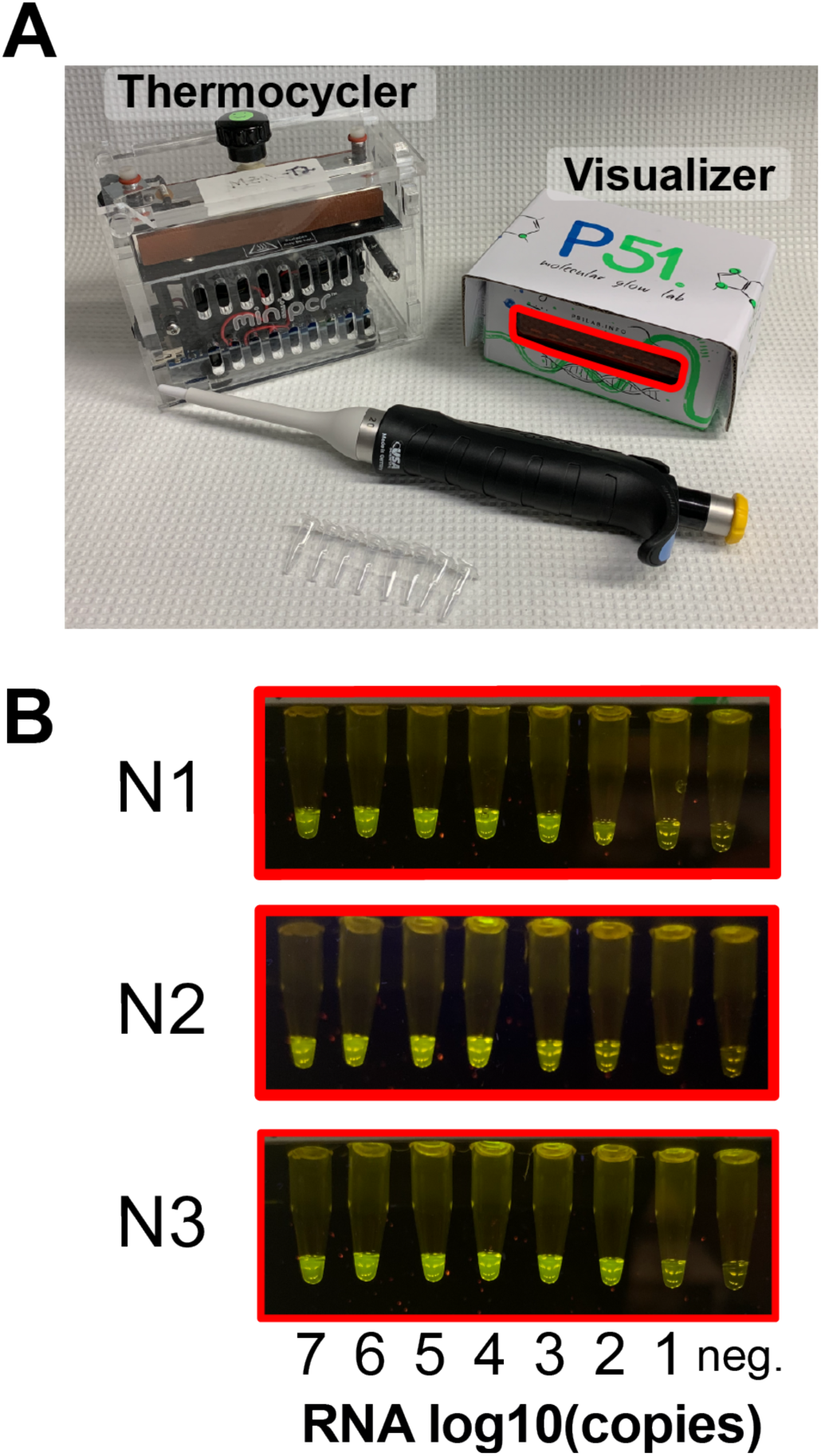
Detection of SARS-CoV-2 RNA using CREST. (A) The miniPCR mini16 thermocycler and P51 molecular fluorescence visualizer used in this study. Both are portable, can be operated with batteries and have minimal footprint. (B) Fluorescence visualization of N1, N2, and N3 synthetic targets using P51 visualizer.

To reduce the second hurdle, the visualization of test outcomes, we explored options available for the detection of Cas13 activity. When bound to its target Cas13 catalyzes the non-specific cleavage of RNAs. This target-specific recognition can be detected in many ways, for example either by lateral flow immunochromatography or by fluorescence visualization through the use of a fluorescein and quencher conjugated poly-U RNA cleavage reporter. Lateral flow test strips are a promising detection method. These strips utilize capillary action to move analytes through a solid support material striped with antibodies that can detect them, resulting in binary readouts. While simple to use and read—they may eventually enable in-home testing—their current availability is limited and they are expensive relative to the cost-per-test (Fig. 3B, Sup. Spreadsheet). For this reason, we sought an affordable, scalable, easy-to-interpret, solution for visualization of positive results. In the CREST protocol, we use a P51 cardboard fluorescence visualizer, powered by a 9V battery, for the detection of Cas13 activity instead of immunochromatography (Fig. 2).

**Figure 3.**
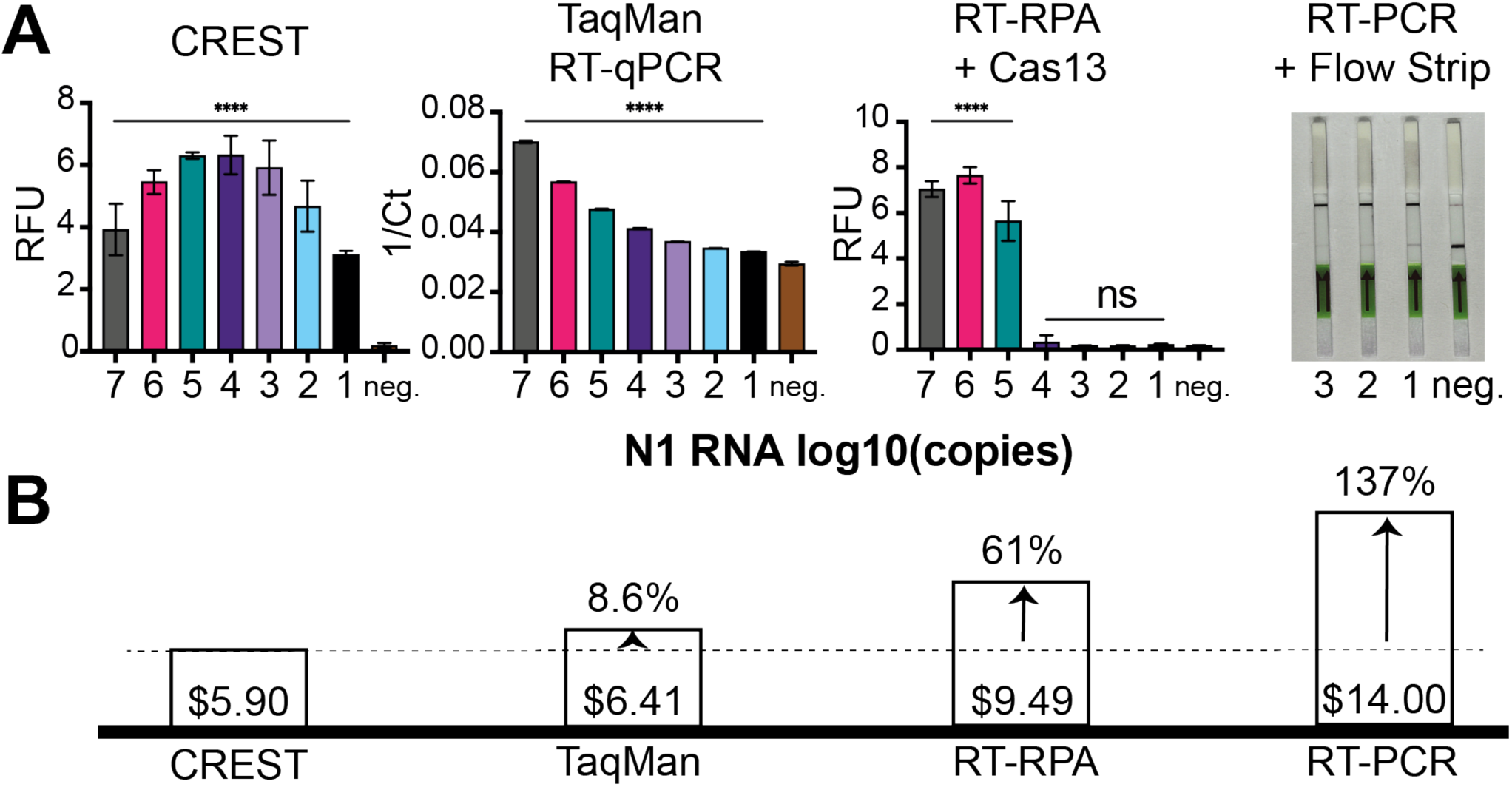
Comparative analysis of method sensitivity and reagent cost per test. (A) Comparison of method sensitivity using a quantitative fluorescence detection instrument. From left to right CREST, TaqPath™ 1-Step RT-qPCR and RT-RPA + Cas13a detection. Far right, RT-PCR + Cas13 detection visualized with lateral flow strips. (B) Associated costs of reagents per test of each testing method (excluding upfront instrumentation costs). A test is defined as a single sample run in triplicate.

To validate a streamlined workflow with these devices, we measured the presence of the three viral sequences that correspond to the CDC RT-qPCR test^19^. Briefly, we PAGE purified annealed synthetic DNA oligonucleotides flanked by an upstream T7 RNA polymerase promoter that encode sequences corresponding to the N1, N2, and N3 sites in the SARS-CoV-2 nucleocapsid gene (Table S3). Next, we transcribed the DNA *in vitro* to obtain target RNAs. After Cas13 purification and extensive optimization of the reaction conditions (Fig. S1, S2), we used these targets to determine the detection limit of the CREST protocol and found that we could detect up to 10 copies of a target RNA molecule per microliter (Fig. 2B). This result shows that CREST is as sensitive as the corresponding RT-qPCR in our hands, demonstrating the power of CREST for pairing a thermal cycling amplification step (PCR) with a linear amplification step (transcription), combined with enzymatic signal amplification through fluorescence detection.

Next, we quantitively compared CREST’s sensitivity and cost to established methods. First, we compared it to RT-qPCR (one-step TaqMan assay) and found that, while being similarly sensitive, CREST’s reagents cost less than RT-qPCR’s even at the low-scale of our pilot experiment (Fig. 3, Sup. Spreadsheet). In addition, the upfront cost of CREST instrumentation is 30 to 50 times lower. Second, we compared the RT-PCR amplification step of CREST to Cas13-based protocols that utilize RT-Recombinase Polymerase Amplification (RT-RPA). We found thermal cycling amplification (20 cycles) to be substantially more efficient with comparable amplification reaction times (Fig. 3A, Fig. S3). Moreover, in stark contrast to the proprietary, high-cost, relatively small batch production rates of reagents required for RPA, Taq polymerase, which has been in high-volume production for decades, and has been a workhorse of modern molecular biology, is readily accessible, stable at room temperature, and lowers costs significantly (Fig. 3B). Lastly, we compared lateral flow test strip visualization to CREST, and while we found them to equally as sensitive as fluorescent detection methods (Fig. 3A), their high cost and difficulty to obtain can limit their distribution and scaled use in this pandemic.

To test the efficacy of CREST on human samples, we obtained 64 de-identified nasopharyngeal swabs from individuals collected from by the Santa Barbara County Department of Public Health. We purified RNA from these samples, which were stored in viral transport media, using a commercially available RNA extraction kit (QIAamp Mini Elute Virus Spin Kit, Qiagen). We used this RNA as input for a parallel comparison between CREST and the CDC-recommended one-step TaqMan assay (our end-to-end CREST protocol is included in the Supplemental Information). Considering that CREST was designed to provide a binary outcome, we fit the CREST-to-TaqMan comparison to a sigmoid. We then calculated a goodness-of-fit R-value between both assays for detection of N1, N2, and RNaseP in 14 samples which contained an unspecified, non-zero number of confirmed positive cases (Fig. 4). These analyses revealed high concordance between CREST and TaqMan assays (R^2^ > 0.9 for SARS-CoV-2 genes and R^2^ > 0.79 for RNaseP). Of note, CREST appears to be more sensitive than TaqMan in its detection of N1 whereas the converse appears to be true for N2. Additionally, we conducted a similarly structured comparative pilot surveillance study on asymptomatic individuals where oropharyngeal self-sampling was used to obtain 95 samples and no positives were detected (Fig. S5).

**Figure 4.**
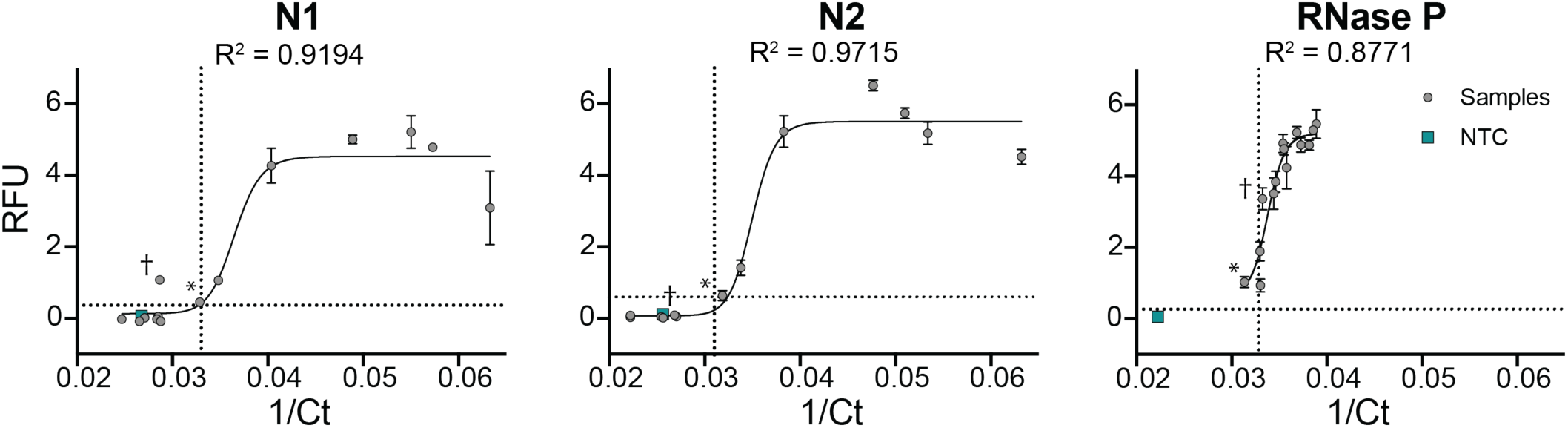
Comparative analysis between CREST and TaqMan for SARS-CoV-2 N1 and N2 sites and RNaseP. De-identified human samples are shown as grey circles. Error bars: Standard deviation of N = 4. Dotted lines indicate the detection threshold for each assay.

Finally, while we designed CREST to be an accessible and scalable assay for detecting SARS-CoV-2, it still requires RNA extraction using commercial kits, which limits its widespread adoption. In a companion paper, we present a method called PEARL (Precipitation Enhanced Analyte RetrievaL) which uses common laboratory reagents to bypass this limitation^29^. To lift the final obstacle to CREST’s accessibility, we coupled PEARL to CREST and found that commercial RNA extraction could be omitted (Fig. S6).

## Discussion

The unparalleled spread of SARS-CoV-2 demands orthogonal solutions to expand testing capacities. Here, we demonstrate an optimized Cas13-based protocol that uses accessible reagents and affordable equipment, while maintaining sensitivity that is comparable to established testing methods. Considering the need for a rugged, equitable, and scalable test, we couple spartan, Bluetooth-enabled thermocyclers that can be plugged into a battery and run using mobile device applications, with simple plastic-filter-based, LED visualizers. Results can be simply documented with a smartphone camera and uploaded to the cloud, enabling distributed point-of-care (POC) testing. Operational costs could be further lowered by multiplexing the amplification of viral targets.

Current methods to detect SARS-CoV-2 presence or prior exposure in patient samples rely on detection of the viral genome or antibodies in nucleic acid and serological tests, respectively^24^. The detection of viral genome sequences by RT-qPCR is recommended by public health organizations worldwide. However, these recommendations fail to account for the massive need for widespread testing. Implementing this method alone places a single large burden on the manufacturing capacity for RT-qPCR testing reagents and consumables, specialized instrumentation, and trained laboratory personnel, thus hampering widespread testing. Other POC diagnostic platforms still require specialized instrumentation and are low throughput (e.g. Abbott ID NOW).

These limitations underscore the need for alternative methods, such as CRISPR-based detection, to relieve some of the strain on the global supply chain for testing reagents. These state-of-the-art methods are scalable, do not entail specialized instrumentation, and require very little specialized training. We designed CREST to capitalize on the strengths of both RT-qPCR and CRISPR-based detection, while addressing their shortcomings. CREST exploits the robustness and reliability of PCR, while harnessing the combined sensitivity and convenience of a coupled transcription-detection reaction. CREST can be run, from RNA sample to result, with no need for AC power or a dedicated facility, with minimal handling in ∼2 hours. Because the signal saturates even at low input levels (Fig. 2B) CREST offers the added advantage of binary result interpretation, similar to lateral flow test strips, but at a fraction of the price, as it does not require the costly antibodies or antibody-conjugates needed for lateral flow immunochromatography. Finally, while purified Cas13 is not yet commercially available, we were able to purify enough protein from a 1-liter bacterial culture for more than 500,000 reactions, and we can provide aliquots of protein upon request. Increasing the potential for field deployment of Cas13 for massive POC testing will require additional optimizations, such as lyophilization. Indeed, our Cas13 master mix is not affected significantly by undergoing numerous freeze-thaw cycles (Fig. S4), which increases CREST’s potential for field deployment by overcoming the need for professional laboratories.

Regular testing may be a key determinant impacting the ability to ascertain foci of infection and reduce false negative rates. Recent studies document that in many individuals, even those with active symptoms, initial tests can be negative, while subsequent tests are positive, and intermittent detection of the virus in samples can also occur^25–28^. Despite the biochemical reliability of currently-available diagnostics, the initial quality of a sample and the viral load in the patient can affect the likelihood of a false result. Because of its low costs and ease of use, CREST could be employed for regular testing as well as for disambiguation of results obtained with established methods.

## Acknowledgements

We thank Feng Zhang and Michael Springer for the kind gift of Cas13a purification plasmids. We thank Radu Gogoana for his helpful comments on scalability and Cameron Myhrvold for his thoughts on assay design. We thank Sebastian Kraves and Ezequiel Alvarez-Saavedra for the P51 visualization devices. We are grateful to Stewart W. Comer of the Santa Barbara County Public Health Department for testing advice and for granting access to SARS-CoV-2 positive samples. We thank Keith Blaha for designing a web database to store CREST data. We thank UC Santa Barbara Office of Research and IEX for their generous support. We thank Jennifer Smith and the Biological Nanostructures Laboratory within the California NanoSystems Institute for their equipment. We thank Monte Radeke for use of his FPLC, Apeel Sciences for use of their FPLC columns and Kevin Brackett for their technical assistance. Finally, we thank all essential workers for keeping society running. Without them this work would have never happened.

