## Supplemental Information and Methods for "A Scalable, Easy-to-Deploy, Protocol for Cas13-Based Detection of SARS-CoV-2 Genetic Material"

---

### SUPPLEMENTAL INFORMATION

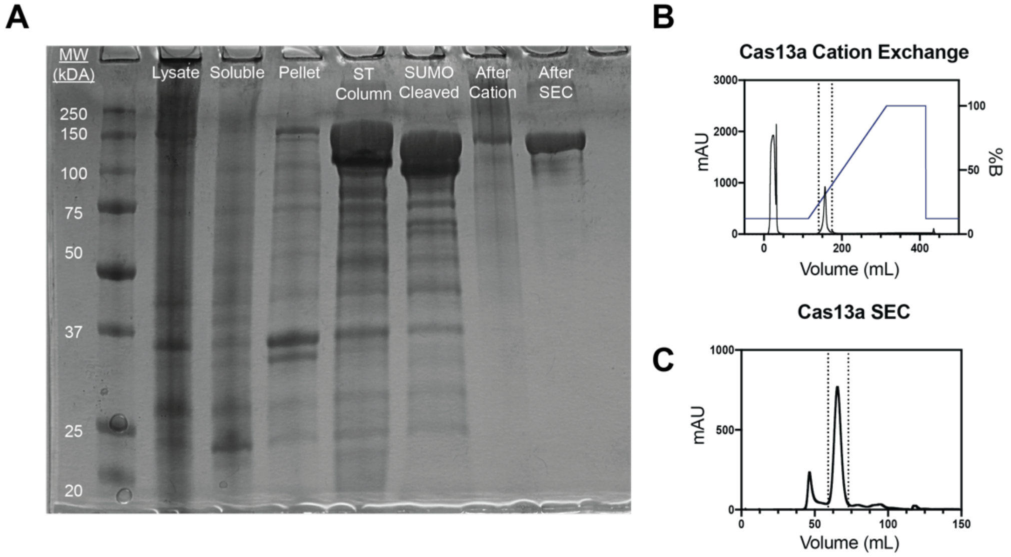

**Supplemental Figure 1.** Purification of Cas13a. (A) SDS-PAGE gel of steps in Cas13a purification. ST column = Strep-Tactin column. (B) Representative trace for cation exchange column. Dashed lines indicated the cut-off for fraction pooling. (C) Representative trace for size exclusion column (SEC). Dashed lines indicated the cut-off for fraction pooling.

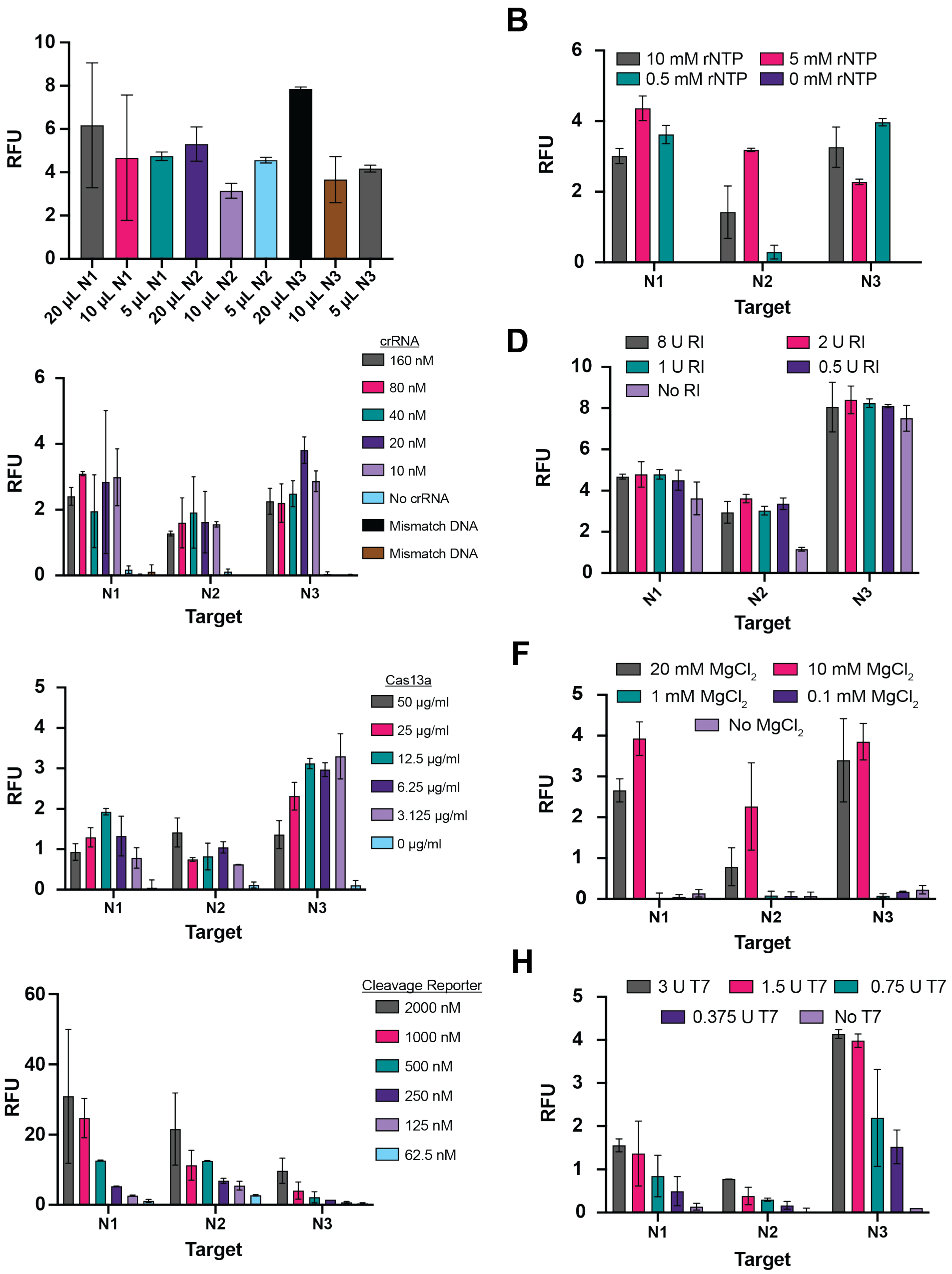

**Supplemental Figure 2.** Optimization of Cas13a detection parameters. (A) The total volume of Cas13a detection reaction does not impact signal sensitivity (B) Cas13a detection under various rNTP final concentrations (C) Cas13a detection with different amounts of crRNA, Mismatch DNA = N2 and N3 synthetic DNA for N1 target, etc. (D) Cas13a detection with various final concentrations of murine RNase Inhibitor (RI). (E) Cas13a detection with various Cas13a final concentrations. (F) Cas13a detection with various $MgCl_2$ final concentrations. (G) Cas13a detection with various cleavage reporter concentrations. (H) Cas13a detection with various T7 RNA polymerase concentrations. All experiments were performed in triplicates and results shown are baseline corrected fluorescent values after 30 minutes of incubation at 37°C (error bars = SD).

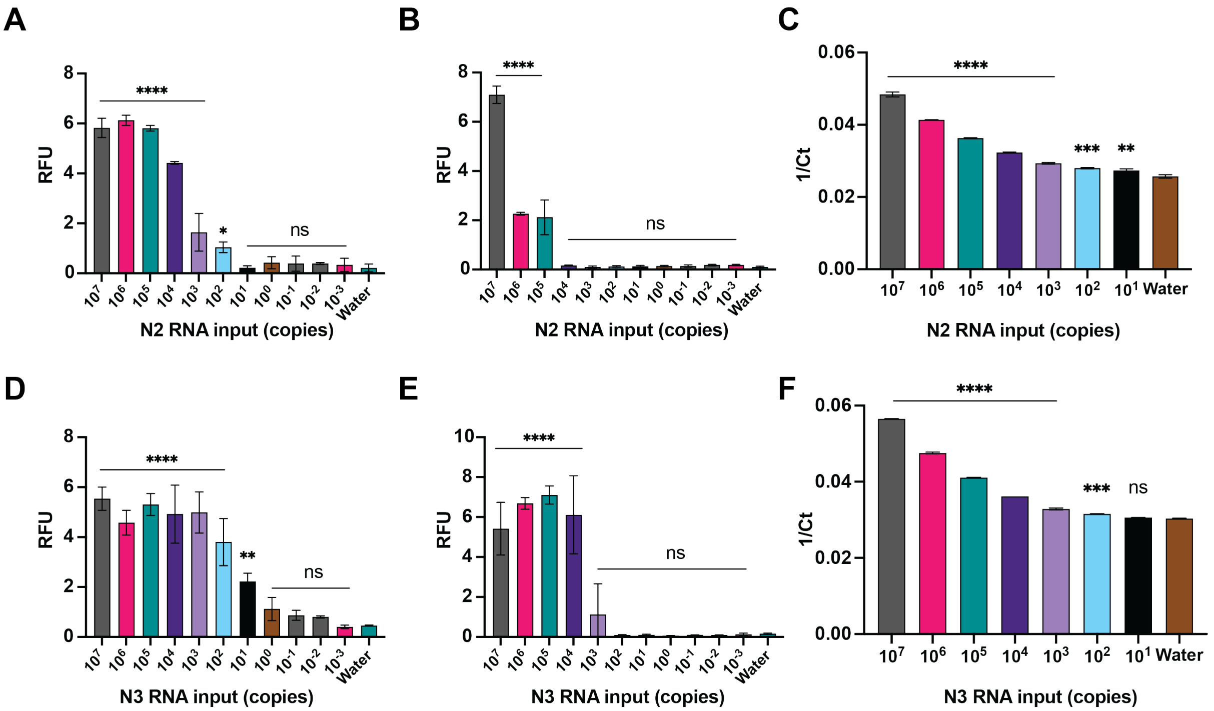

**Supplemental Figure 3.** Detection sensitivity for N2 and N3 targets. (A & D) RT-PCR followed by Cas13a detection for N2 and N3 targets. (B & E) RT-RPA followed by Cas13a detection for N2 and N3 targets. (C & F) TaqMan RT-qPCR for N2 and N3 targets. Experiments were performed in duplicates or in triplicates (error bars = SD). Statistical significance was determined using one-way ANOVA with Dunnett's method (relative to water negative control).

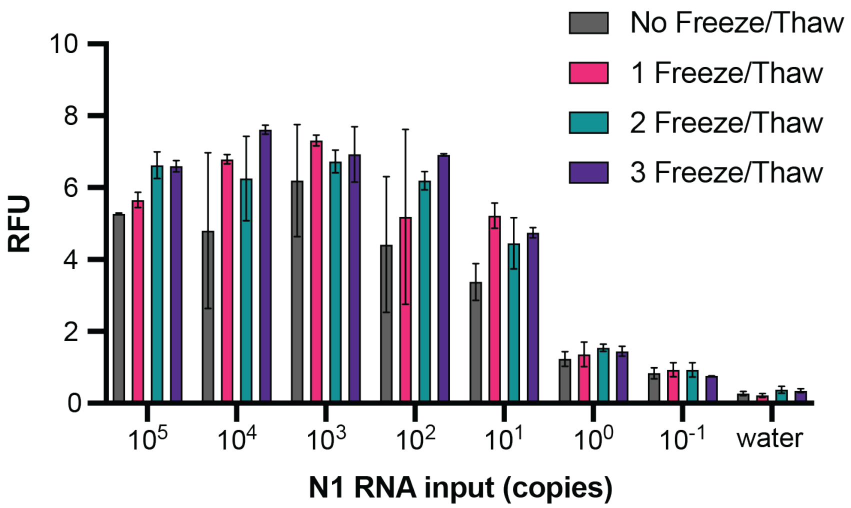

**Supplemental Figure 4.** Cas13 detection master mix shows no loss in sensitivity of SARS-Cov-2 N1 upon multiple freeze-thaw cycles. Experiments were performed in duplicate (error bars = SD).

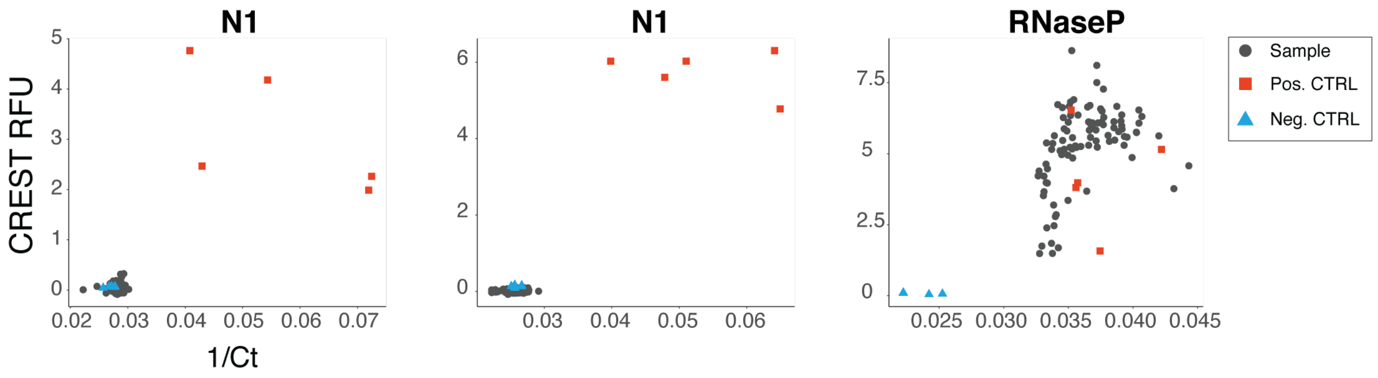

**Supplemental Figure 5.** Side-by-side comparison of CREST and RT-qPCR done on 95 de-identified samples taken from asymptomatic individuals for N1, N2 and RNaseP with positive and negative control samples.

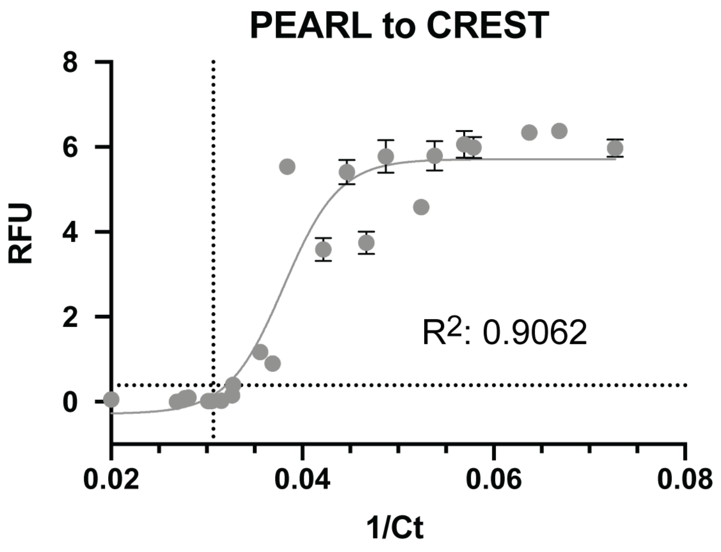

**Supplemental Figure 6.** Demonstration of coupling PEARL, column-free, RNA extraction method to both CREST (Y-axis) and RT-qPCR (X-axis) for 25 independent clinical samples. Dotted line indicates limit of detection for each assay. Experiments were performed in duplicate (error bars = SD).

### METHODS

**Table 1.** PCR and RPA primer sequences. The minimal T7 RNA polymerase promoter is indicated in uppercase. We added a *gggc* spacer after the minimal promoter sequence to increase transcription rates and prevent transcript heterogeneity in the 5' leader.

| Name | Sequence 5' to 3' |
|---|---|
| N1 fwd | *gaaat*TAATACGACTCACTATA*gggc*gaccccaaaatcagcgaaat |
| N1 rev | tctggttactgccagttgaatctg |
| N2 fwd | *gaaat*TAATACGACTCACTATA*gggc*ttacaaacattggccgcaaa |
| N2 rev | gcgcgacattccgaagaa |
| N3 fwd | *gaaat*TAATACGACTCACTATA*gggc*gggagccttgaatacaccaaaa |
| N3 rev | Tgtagcacgattgcagcattg |
| RNase P fwd | gaaatTAATACGACTCACTATAgggagatttggacctgcgagcg |
| RNase P rev | gtgagcggctgtctccacaa |

**Table 2.** Synthetic gRNA sequences. Shared Cas13 crRNA denoted in uppercase.

| Name | Sequence 5' to 3' |
|---|---|
| N1 gRNA | GAUUUAGACUACCCCAAAAACGAAGGGGACUAAAACagggucccaccaaacguaaugcggggugc |
| N2 gRNA | GAUUUAGACUACCCCAAAAACGAAGGGGACUAAAACgcugaagcgcugggggcaaauugugcaa |
| N3 gRNA | GAUUUAGACUACCCCAAAAACGAAGGGGACUAAAACuagcaggauugcgggugccaaugugauc |
| RNase P gRNA | GAUUUAGACUACCCCAAAAACGAAGGGGACUAAAACguccgcgcagagccuucaggucagaacc |

**Table 3.** Template DNA for *in vitro* transcription of RNA targets. The minimal T7 RNA polymerase promoter is indicated in uppercase.

| Name | Sequence 5' to 3' |
|---|---|
| synTarget N1 | gaaatTAATACGACTCACTATAggggaccccaaaatcagcgaaatgcaccccgcattacgtttggtggaccctcagattcaactggcagtaaccaga |
| synTarget N2 | gaaatTAATACGACTCACTATAgggttacaaacattggccgcaaattgcacaatttgcccccagcgcttcagcgttcttcggaatgtcgcgc |
| synTarget N3 | gaaatTAATACGACTCACTATAggggggagccttgaatacaccaaaagatcacattggcacccgcaatcctgctaacaatgctgcaatcgtgctaca |

**Table 4.** RNA cleavage reporter sequences.

| Name | Sequence 5' to 3' |
|---|---|
| CREST Fluorescent Cleavage reporter | 6-FAM (Fluorescein) – $(U)_7$ – Iowa Black FQ |

| | |
|---|---|
| Lateral Flow Strip Cleavage reporter | 6-FAM (Fluorescein) – $(U)_{14}$ – Biotin |

#### *In vitro* transcription of synthetic targets

Single strand DNA ultramers were purchased from Eton Biosciences (San Diego, CA, USA). The ssDNA ultramers were dissolved in nuclease free water at a concentration of 10 µg/µL. To generate dsDNA templates, the ultramers were annealed in 2X annealing buffer (100 mM potassium acetate, 30 mM HEPES-KOH pH 7.4, 2 mM magnesium acetate) by heating at 95˚C for ten minutes in an aluminum heat block. While remaining in the heat block, the annealed ultramers were slowly cooled to room temperature. The annealed ultramers were subsequently separated on 4-20% gradient TBE PAGE gels (Thermo Fisher Scientific) at 150V. Gels were stained with SYBR-SAFE (Thermo Fisher Scientific) diluted 1:10,000 in 1X TBE buffer for 10 minutes, and the annealed DNA were excised from the gel using a clean razor blade while being visualized with a transilluminator. Gel fragments were placed in 1.5 mL tubes with 500 µL TE buffer (10 mM Tris pH 8.5, 1 mM EDTA) and shaken overnight at 700 rpm, 16˚C. TE buffer containing the eluted DNA templates was removed from the gel fragment and placed into a fresh tube. The annealed DNA templates were subsequently precipitated by mixing with one volume of ice-cold isopropanol and one ninth volume of 3 M sodium acetate pH 5.5, incubation at -80˚C for thirty minutes, and collected via centrifugation at 20,000 x *g* (4˚C) for thirty minutes. Pellets were washed with 1 mL ice-cold 80% ethanol and air-dried for ten minutes. The DNA was resuspended in 10 µL of nuclease free water. dsDNA quantity and purity were assessed using a NanoDrop ONE instrument (Thermo Fisher Scientific).

DNA templates were transcribed using the Megascript T7 Transcription kit (Thermo Fisher Scientific) 500 ng of PAGE-purified dsDNA were used as input for each reaction, all other reagents were added according to the manufacturer's protocol. Transcription was allowed to proceed overnight at 37˚C. After transcription, DNA templates were removed by incubating the reaction mixture with 2 units of DNase (as detailed in the Megascript T7 Transcription kit) at 37˚C for 15 minutes. Ammonium acetate was added to inactivate the DNases according to the manufacturer's protocol.

#### Purification of synthetic RNA targets

Synthetic RNA targets were purified by phenol extraction. Briefly, the RNA was subjected to two sequential extractions, the first using one volume of acid phenol (5:1 phenol:chloroform, pH 4.5, Thermo Fisher Scientific) and a second one using one volume of chloroform. After addition of the organic solvents, the samples were mixed by vortex and centrifuged at 12,500 x *g* (4˚C) for 15 minutes. The aqueous phase was recovered each time. After the second extraction, the RNA-containing aqueous phase was mixed with one volume of ice-cold isopropanol and one ninth volume of 3M sodium acetate pH 5.5, incubated at -80˚C for 20 minutes and the precipitated RNA was collected by centrifugation at 16,000 x *g* (4˚C) for 20 minutes. The RNA pellets were washed with 1 mL of ice-cold 80% ethanol and air dried for 10 minutes. The RNA was resuspended in 10 µL of nuclease free water. RNA quantity and quality were assessed using a NanoDrop ONE instrument (Thermo Fisher Scientific).

#### Reverse transcription and cDNA Amplification

Reverse transcription (RT) was performed using RevertAid Reverse Transcriptase (200 U/µL, Thermo Fisher Scientific) in presence of murine RNase inhibitor (New England Biolabs). Reaction components were mixed in 0.2 mL tubes as follows:

| | |
|---|---|
| RNA template | 5 µL |
| Gene Specific Primer Mix (5 µM each primer) | 1 µL |
| 5X Reaction Buffer | 2 µL |
| dNTPs (10 µM each NTP) | 1 µL |
| RevertAid Reverse Transcriptase | 0.5 µL |
| Murine RNase inhibitor (40 U/ µL) | 0.5 µL |
| **TOTAL VOLUME** | 10 µL |

All components were gently mixed by pipetting and the reactions were collected by centrifugation using a tabletop centrifuge. The reaction mixture was then heated to 42°C for 30 minutes in a heat block. Upon completion, reaction was immediately placed on ice. The resulting cDNAs were used as input for two amplification methods; polymerase chain reaction (PCR) and recombinase polymerase amplification (RPA).

#### PCR amplification

Target DNA molecules were amplified by PCR using a *Taq* DNA polymerase master mix (New England Biolabs). Reaction components were mixed in 0.2 mL tubes as follows:

| | |
|---|---|
| Nuclease Free Water | 17 µL |
| 5X Master Mix | 5 µL |
| Gene Specific Primer Mix | 1 µL |
| RT Product | 2 µL |
| **TOTAL VOLUME** | 25 µL |

The reactions were mixed, and target DNAs were amplified in a mini16 thermocycler (miniPCR bio, Cambridge, MA) using the following thermal profile:

| | |
|---|---|
| 98°C | 2 min |

| | | |
|---|---|---|
| 98°C | 15 sec | 20X |
| 60°C | 15 sec | |
| 72°C | 15 sec | |
| 72°C | 5 min | |

**RPA amplification**

Target DNA molecules were amplified by RPA using TwistAmp Liquid Basic RPA kits (TwistDx, Maidenhead, UK). The RPA master mix was assembled as follows:

| Component | Volume per reaction |
|---|---|
| 2X Buffer | 10 μL |
| dNTPs (10 μM each NTP) | 3.7 μL |
| 10X Basic E Mix | 2 μL |
| Primer Mix (5 μM each primer) | 2 μL |
| Vortex and centrifuge | |
| 20X Core reaction Mixture | 1 μL |

18 μL aliquots of the RPA master mix were added to 0.2 mL tubes and 1 μL of kit-supplied magnesium acetate was added to each reaction. Finally, the reactions were started by addition of 1 μL of cDNA and incubation at 42°C for 20 to 40 minutes.

**LwaCas13a purification**

Plasmids expressing Cas13a were kind gifts from Michael Springer and Feng Zhang (pC013: Addgene #90097). Purification of Cas13a was performed according to published protocols (Kellner et al. Nature Protocol 2019). Briefly, Rosetta 2(DE3)pLysS cells (Millipore) harboring the Twinstrep-SUMO-huLwaCas13a were grown in TB medium until an OD600 of 0.6 was achieved. Protein expression was induced by treating the cells with 250 μM IPTG overnight at 16°C. Cells were harvested and lysed in lysis buffer (20 mM Tris pH 8.0, 500 mM NaCl, 1 mM DTT, + cOmplete$^{TM}$ protease inhibitor tablets) via sonication. The soluble fraction was incubated with Strep-Tactin agarose (EMD) for 2 hours at 4°C and the non-bound proteins were washed away. Overnight cleavage (4°C) of the SUMO-tag was performed on resin using 50 U of SUMO protease (Thermo Fisher Scientific) and 0.15% NP-40. The next day, Cas13a was eluted from the resin, diluted to 24 mM NaCl, and loaded onto a cation exchange column (GE Lifesciences) equilibrated in 20 mM Tris pH 7.5, 24 mM NaCl, 5% Glycerol, 1 mM DTT. Fractions containing Cas13a were pooled, concentrated, and loaded onto an SEC200 column (GE Lifesciences) equilibrated in 50 mM Tris pH 7.5, 600 mM NaCl, 2 mM DTT. Fractions containing Cas13a were pooled and buffer exchanged into Cas13a storage buffer (50 mM Tris pH 7.5, 600 mM NaCl, 5% Glycerol, 2 mM DTT).

Protein concentration was determined by absorbance at 280 nm. Typical yields were 7 mg/L.

#### Cas13a-based Detection

Cas13a was used for site-specific detection with three readouts; fluorescent detection using a Quantstudio5 qPCR instrument (Applied Biosystems), or using a P51 visualizer (miniPCR bio), and visual detection with lateral flow strips. Fluorescent readouts followed similar protocols, and lateral flow strips utilized a different cleavage reporter that was 3' biotinylated instead of the chemical addition of Iowa Black Quencher. Cas13a detection was performed in Cas13a cleavage buffer (1x: 40 mM Tris pH 7.5, 1 mM DTT) supplemented with 1 mM rNTPs (Thermo Fisher Scientific), 2 U/μL RNase Inhibitor (New England Biolabs), 0.125 μM cleavage reporter (Integrated DNA Technologies), 1.5 U/μL T7 RNA Polymerase (Lucigen), 6.3 ng/μL LwaCas13a, 20 nM Cas13 crRNA and 9 mM $MgCl_2$ unless otherwise indicated. Reactions were typically composed of 4 μL Cas13a cleavage solution and 1 μL of sample (DNA standard, RT-RPA or RT-PCR product) in a single well of a 384 well-plate. Plates were sealed and florescence was acquired every 5 minutes for 30 minutes at 37°C in a Quantstudio5 qPCR instrument. An initial reading was taken at time = 0 and subtracted from results shown. Experiments were performed in triplicates. Alternatively, reactions were placed in PCR tubes and incubated at 37°C for 5-30 minutes and fluorescence was detected visually using the P51. We not that visual detection should be done blinded and that accuracy gets better after practice and in a dark room. For lateral flow strips, the Cas13a reaction mixture was scaled to 20 μL, incubated at 37°C for 0.5-1.5 hours and read via lateral flow detection (Metsky et al. Biorxiv 2020).

#### RT-qPCR, Taqman Assay (CDC recommended protocol)

TaqPath 1-step RT-qPCR MasterMix (Cat #pdt A15300) was purchased from Applied Biosystems. A master mix of was prepared using the established CDC protocol (https://www.fda.gov/media/134922/download). 15 μL of the master mix were dispensed into the qPCR plate before addition of input RNA. Serial dilutions of *in vitro* transcribed RNA, ranging from 200 fg/μL to 2 ag/μL and corresponding to $10^7$ to $10^2$ viral copies/μL, were prepared in nuclease free water. 5 μL of each RNA target dilution was added to the wells containing the corresponding TaqMan primers and probes. For no template controls, 5 μL of nuclease free water was used in place of template RNA. The plate was sealed with film and spun down by a pulse in a centrifuge. All samples were run in technical duplicates. The reactions were run on a real time PCR instrument (BioRad CFX96 Touch) using the following thermal profile:

| | |
|---|---|
| 25°C | 2 min |
| 50°C | 15 min |
| 95°C | 2 min |

| | | |
|---|---|---|
| 95°C | 15 sec | 44X |
| 55°C | 60 sec | |
| | Plate Read | |
| 4°C | hold | |

Data were analyzed using the BioRad CFX Maestro software. Thresholds for Cq value determination were obtained by regression after baseline subtraction.

#### Statistical Analysis

All statistical analyses were performed using Prism v8.0. To determine statistical significance, a one-way ANOVA with Dunnett's multiple comparison was performed.

#### Extraction of RNA by PEARL

De-identified human buccal swabs (1 mL) in viral transport media were supplemented with 4 µL of the heat-inactivated SARS-CoV-2 virus (ATCC® VR-1986HK™). Aliquots of 250 µL were mixed in a 1:2 (v/v) sample:PEARL lysis solution (0.5% IGEPAL CA-630, 450 mM sodium acetate, 20 % glycerol, 20 mM TCEP, 50 µg/ml linear polyacrylamide, and 20 mM HEPES-KOH pH 7.2) ratio and incubated for 5-minutes at room temperature. Next, RNA was precipitated on ice, for 10 minutes, using one volume of cold isopropanol. The precipitated material was collected by centrifugation at 19,000 x g for 10 minutes, washed once with 75% ethanol, air-dried for 5 minutes at room temperature and solubilized in 20 µl of nuclease-free water. CREST reactions were performed as described above.
